# Strategies for building computing skills to support microbiome analysis: a five-year perspective from the EDAMAME workshop

**DOI:** 10.1101/631267

**Authors:** Ashley Shade, Taylor K. Dunivin, Jinlyung Choi, Tracy K. Teal, Adina C. Howe

**Affiliations:** Department of Microbiology and Molecular Genetics, Michigan State University, East Lansing, MI 48824; Department of Plant, Soil and Microbial Sciences, Michigan State University, East Lansing, MI 48824 USA; Department of Agricultural and Biosystems Engineering, Iowa State University, Ames, IA 50011 USA; Data Carpentry, Davis, CA USA

## Abstract

Here, we report our educational approach and learner evaluations of the first five years of the Explorations in Data Analysis for Metagenomic Advances in Microbial Ecology (EDAMAME) workshop, held annually at Michigan State University’s Kellogg Biological Station from 2014-2018. We hope this information will be useful for others who want to organize computing-intensive workshops and encourage quantitative skill development among microbiologists.

**Importance:** High-throughput sequencing and related statistical and bioinformatic analyses have become routine in microbiology in the past decade, but there are few formal training opportunities to develop these skills. A week-long workshop can offer sufficient time for novices to become introduced to best computing practices and common workflows in sequence analysis. We report our experiences in executing such a workshop targeted to professional learners (graduate students, post-doctoral scientists, faculty, and research staff).

## Introduction

Over the last decade, two important advances have fostered a new era in biomedical and environmental research. First, it is now recognized that microbial communities (“microbiomes”) play essential roles for the health of the environments and the hosts that they inhabit. Second, advances in high-throughput sequencing technologies allow observations of the diversity and functional potential of microbiomes in their habitats (1), captured with spatially and temporally ambitious study designs (2). Together, these advances in knowledge and methodology deepen and broaden our understanding of the centrality of microbiomes for host and environmental health. Because of the economy and accessibility of high-throughput sequencing, researchers can now investigate the diversity of interesting microbiomes and can begin to untangle how this diversity contributes to host or ecosystem health. Efforts to capitalize on the promise of microbiome sequencing data have resulted in information-rich genomic datasets that must be analyzed to gain knowledge of their intricate relationships.

We realized that there was a need for broad computational training in microbiome analysis. In 2014, we were encouraged by Dr. C. Titus Brown (now at University of California-Davis) to offer a microbiome analysis workshop. At the time, he led the Analyzing Next-Gen Sequencing (ANGUS, https://angus.readthedocs.io/en/2018/index.html) Workshop at Michigan State University’s Kellogg Biological Station (KBS). He noted that some ANGUS learners were particularly interested in microbiome analysis and that there were limited offerings for this training. At the time, there were several short-duration workshops focused on specific tools, such as QIIME(4) and mothur(5), as well as a broader, multi-week course, STAMPS (https://www.mbl.edu/education/courses/stamps/), at the Marine Biological Laboratory in Woods Hole, MA USA. There were few workshops that addressed the needs of learners who wanted more information than what could be covered in a day but also could not commit to spending several weeks away. Thus, we suspected that there was a need for broad and economical training in microbiome analysis, especially in the U.S. Midwest.

In response, we created a one-week intensive course to train biologists (from graduate students to faculty) in microbiome-associated sequencing analysis, from raw sequence handling and quality control to statistical analyses and experimental design. We named the course EDAMAME: Explorations in Data Analysis for Metagenomic Advances in Microbial Ecology. Ashley Shade, at the time a new assistant professor in microbial ecology at the Department of Microbiology and Molecular Genetics at Michigan State University, initiated the workshop and started its content development from her materials from a short workshop she offered while training in her post-doctoral advisor’s lab. Tracy Teal was recruited and brought her array of experience and perspective as a leader in the Software and Data Carpentries workshops, which provide general computing training. In the first year, J. Herr, a post-doc in Shannon Manning’s lab at Michigan State who had Data Carpentry training, contributed to developing and implementing the original content. The instruction team expanded in 2016 to include Adina Howe, who was a new faculty at Iowa State and brought important expertise in untargeted metagenome analysis.

Here, we report a five-year perspective on the EDAMAME workshop. We present EDAMAME’s learning objectives, target audience and admissions, instructional team, learning environment, educational strategy and assessment, and community resources. We discuss results from assessment, lessons learned and an outlook for future microbiome training.

## Results

### EDAMAME learning objectives

EDAMAME’s learning objectives were tailored annually to incorporate learners’ changing interests and changes in tools and technology (**Box 1**). As a consequence, we created and retired tutorials as demands changed. However, each year featured foundational tutorials in computing literacy, state-of-the-science tools for microbiome analyses, ecological statistics, and computing best practices.

**Box 1. Overview of learning objectives for the EDAMAME workshop.**

- Develop working proficiency at the command line and with shell.
- Explain the process of high-throughput sequencing, provide an overview of data-handling (quality control, pre-treatment), and discuss their biases.
- Access computing resources: Transfer data and run analyses on Amazon EC2 and/or a high-performance computing cluster.
- Access and/or create version-controlled code and resources on GitHub.
- Discuss steps in the ecological analyses of microbiomes, including alpha and beta-diversity, ordinations, and resemblance metrics.
- Explore datasets and statistically test hypotheses in R.
- Visualize patterns in microbial communities using R.
- Develop a working proficiency with amplicon sequencing workflows and tools (e.g., QIIME, or mothur, or usearch; (3).
- Develop a working proficiency with shotgun metagenomics workflows.
- Become familiar with publicly accessible microbial sequence databases/repositories (e.g., NCBI, MG-RAST, FunGene) and the tools that they offer for deposition and analyses.
- Identify resources for troubleshooting. This includes: how to ask for and where to find general help online, through peer networks, and from workflow-specific resources (e.g., public tutorials and wikis).

### Target learners and admissions

We targeted our applicant pool towards learners who would benefit most from the training and who we expected would share their developed expertise with others to maximize the reach of the workshop’s training. We accepted applicants who were novice in their analysis skillset and did not have apparent access to other resources to support their skill development. We also aimed to promote diversity in scientific discipline (e.g., human, agricultural, environmental), learner gender and background, research institution (e.g., undergraduate-serving, research university, agencies), geography (with special advertising to learners from the Midwest), and academic level (**Figure 1, Figure 2**). We also strove to provide opportunity to international learners and learners from underrepresented backgrounds. To advertise the course, we used social media (Twitter), our website, and professional networks. We also attempted to reach broader audiences by advertising with international scientific networks, especially Ciencia Puerto Rico in 2014 - 2016.

**Figure 1.**
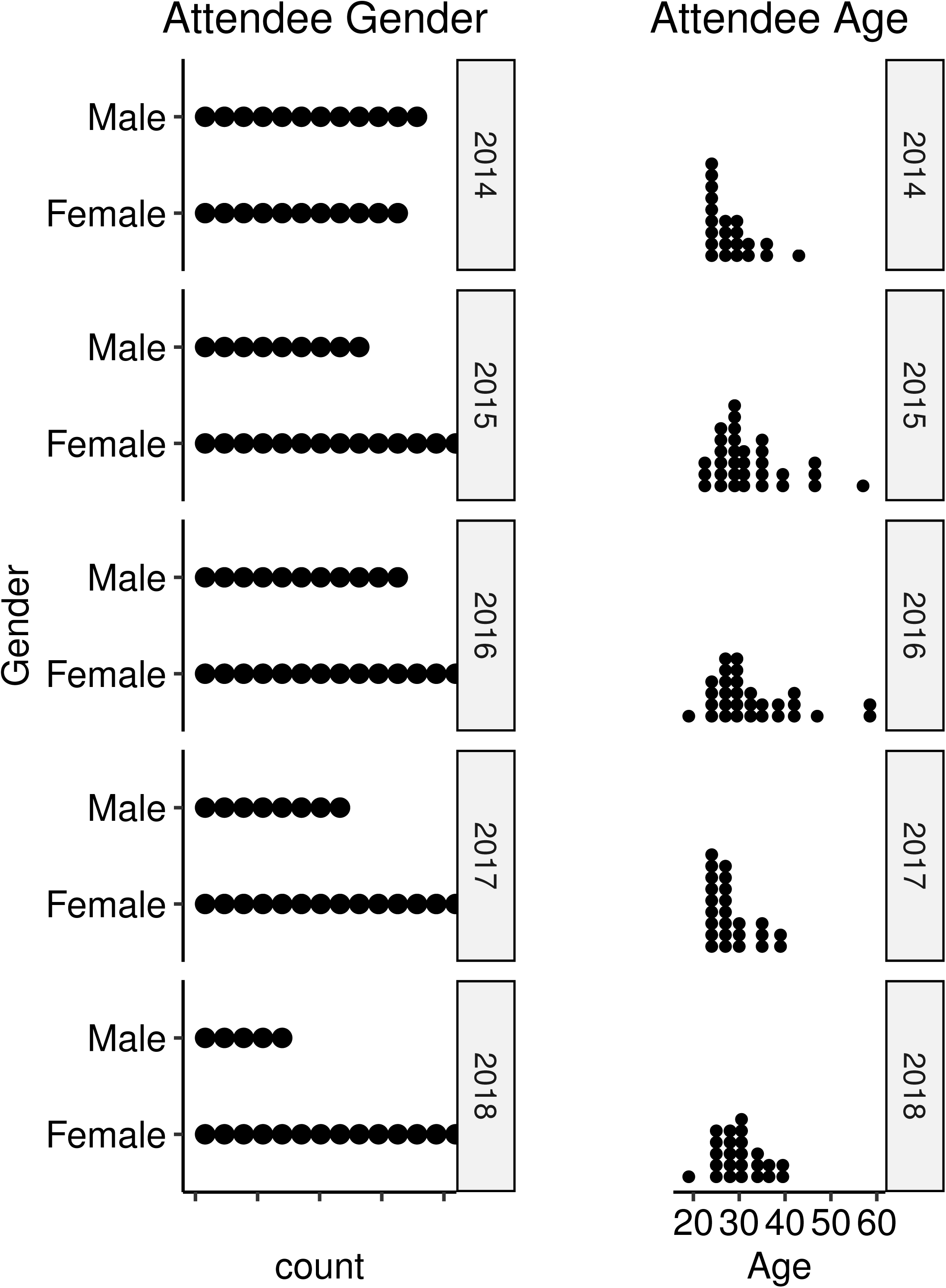
Distributions of EDAMAME learner gender and age, 2014-2018.

**Figure 2.**
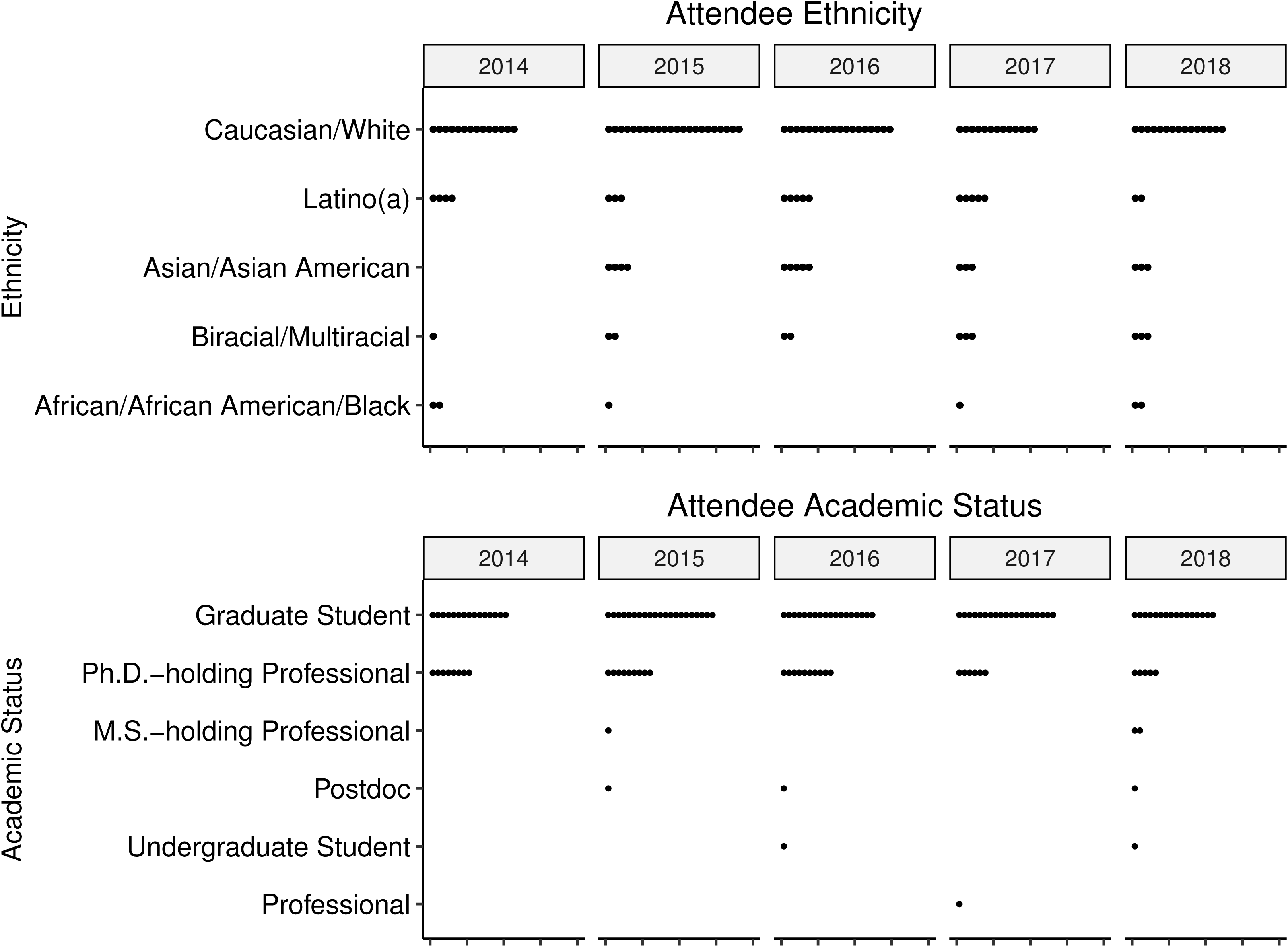
Distributions of EDAMAME learner ethnicity and academic status, 2014-2018.

In each workshop, we could accommodate 23 - 26 learners in the classroom, and applications were oversubscribed every year (**Table 1**). As admissions became increasingly competitive, we began to require (rather than to encourage) that applicants had generated a microbiome dataset prior to the workshop. We found that students who had struggled in an analysis attempt were highly incentivized to maximize their time at the workshop. Also, they could work on their data during office hours and ask specific questions to the instructors and TAs.

**Table 1.** Summary of EDAMAME dates, instructional staff, applicants, and learners from 2014-2018.

| Year | Dates | No. Days | No. TAs | No. Instructors | No. applicants | No. workshop learners<sup>1</sup> |
| --- | --- | --- | --- | --- | --- | --- |
| 2014 | 22 June to 29 June | 8 | 1 | 3 | 50 | 23 |
| 2015 | 21 June to 01 July | 11 | 6<sup>2</sup> | 1 | 93 | 32<sup>3</sup> |
| 2016 | 10 July to 20 July | 11 | 6 | 3 | 62 | 25 |
| 2017 | 06 August to 12 August | 7 | 7 | 3 | 63 | 26 |
| 2018 | 24 June to 30 June | 7 | 10 | 2 | 103 | 26 |

<sup>1</sup>No. workshop learners are from pre- and post- survey responses. Additional local learners participated *ad hoc* and may not have completed surveys.<sup>2</sup> There were two guest TAs in 2015 who participated only in one tutorial each, with the remaining 4 TAs available throughout the workshop. <sup>3</sup>2016 participant data included 3 remote learners who participated in select tutorials.

### Instructional team

A large instructional team was necessary to support EDAMAME’s learning goals. There were one to three lead instructors per year (**Table 1**). The instructors led the course, oversaw admissions, provided lectures and course content, determined guest lectures, and mentored TAs in tutorial development. In the final two years of the workshop, there was also a course coordinator who oversaw conference logistics, fielded learner and applicant questions, and coordinated transportation for learners, guest lecturers, and instructors.

The hands-on nature of the workshop necessitated several dedicated TAs. Multiple instructors and supportive TAs in the classroom allowed us to be immediately responsive to the needs of the learners. TAs led tutorials based on interest and expertise. Having multiple TAs broadened instructional expertise and allowed unscheduled time for each TA to rest when they were not supporting instruction. Most often, new learners struggled with basic syntax and interpreting error messages. Novice TAs (e.g., early graduate students) helped learners trouble shoot common errors, while the more senior TAs and instructors assisted with more complicated hurdles (e.g., software and operating system incompatibilities, experimental design power for data analysis). In addition to instruction, TAs supported the logistical aspects of the course, such as local transportation for learners, purchasing supplies, and assisting learners with unexpected personal needs (e.g., trip to the medical center, forgotten toothbrush). TAs included volunteers (graduate students and post-docs) and graduate assistants partially supported by EDAMAME external funding. Participation in the workshop also offered TAs benefits to engage in teaching opportunities that served diverse audiences.

There were also several invited guest instructors who offered tutorials, technical lectures, and research talks (**Table 2**). Guest instructors varied according to guest availability, learner interests, and workshop duration, but some guest instructors generously provided content every year. Stuart Jones (University of Notre Dame) taught statistical analysis in R; Patrick Schloss and members of his lab (University of Michigan) taught amplion analysis with mothur; Jim Tiedje (Michigan State University) provided a lecture and discussion on the future of microbial ecology. Instructors interacted with the learners during dinner and social time, and this provided an opportunity for learner networking and discussions.

**Table 2.**
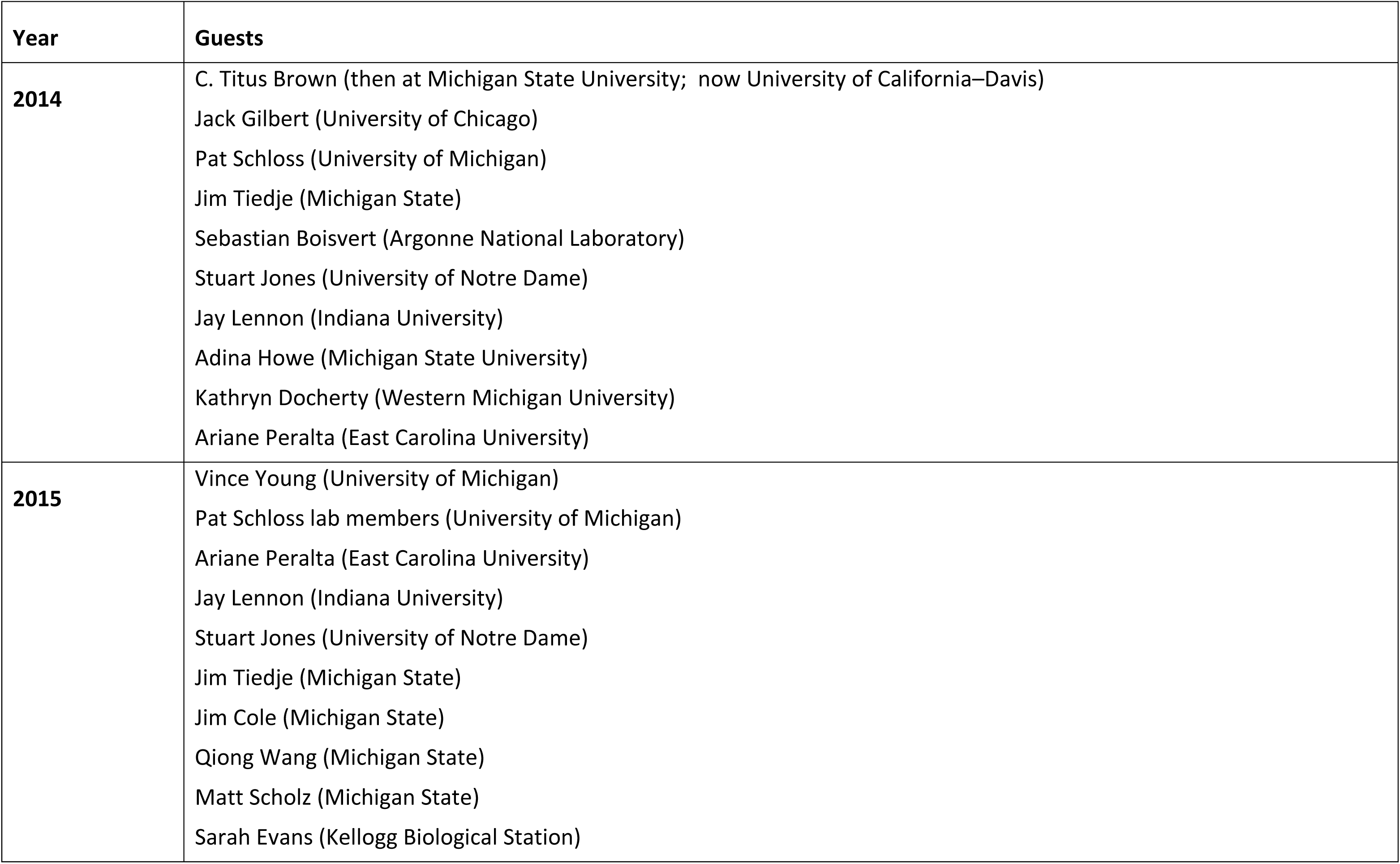
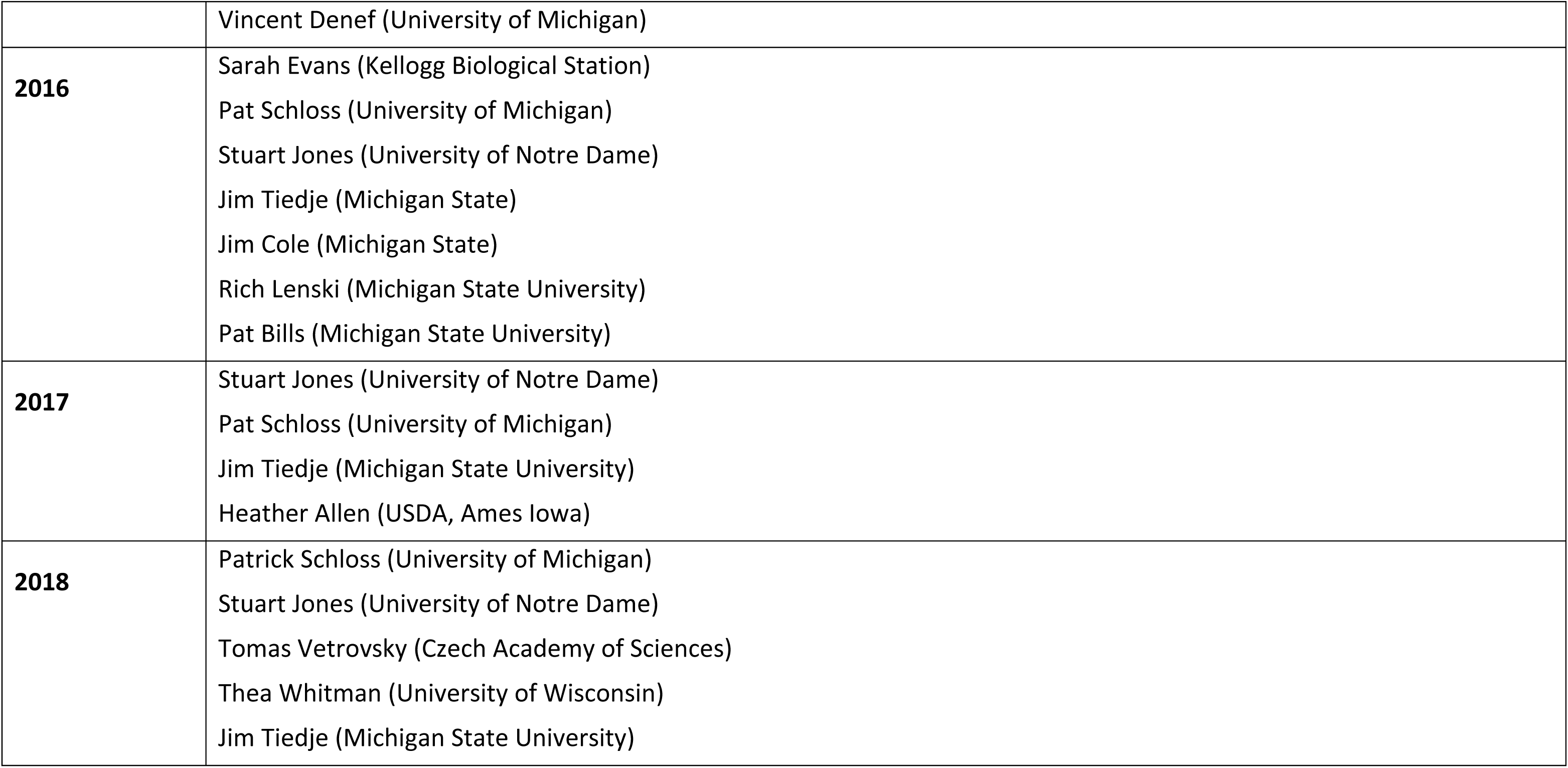
Guest lecturers and instructors for EDAMAME.

### Learning environment and daily schedule

EDAMAME was held at the Kellogg Biological Station (KBS), which offered a remote location, offering an immersive experience for learners and instructors. KBS was also chosen for economy – the room and board rates at KBS were affordable to many (e.g., ∼$370 per week in 2018). Teaching assistants and volunteers provided transportation from the Kalamazoo and Lansing airports to KBS. KBS also provided conference services, dining, wifi, and bonfires. Finally, the natural setting and outdoor activities at KBS provided a respite to time spent in front of the computer.

The length of the workshop varied from 7 - 11 days (**Table 1**), including travel days. The morning schedule included an overview lecture followed by hands-on tutorials and group learning activities. After lunch, we had an afternoon lecture and additional tutorials. We held optional office hours with “choose your own adventure” tutorials and/or lectures on learner-chosen topics during the afternoon break. For example, in 2018 we discussed exact sequence variant analysis. Learners could also ask specific questions about their own data during office hours. After dinner, we held an evening guest lecture in microbiome research. Evenings provided free time for networking and relaxation.

### EDAMAME educational strategy and assessment

EDAMAME’s educational strategy addressed two training needs. First, we offered general training in the fundamentals of introductory computing (*e.g.*, command line, scripting, cloud computing, bioinformatic workflows). This equipped participants with the basic skills needed to independently execute their analyses. We also offered specific training to overcome hurdles particular to microbial metagenomic data analysis and advised on best practices for microbiome analysis. To iteratively assess these strategies, we used a combination of summative and formative assessments to determine participant learning gains.

For the summative assessments, we worked with educational consultants to develop online, anonymous surveys and perform pre-and post-workshop assessments. These assessments evaluated student-reported learning gains and confidence in areas aligned with our learning objectives. The learners created a password to preserve their anonymity while allowing for linking the pre-and post-survey responses. To maximize response rate, we provided dedicated time in the classroom to complete the surveys. The pre-assessment survey was completed on the first full day, and the post-assessment survey was completed on the final day of the workshop. We updated the survey annually to reflect any new or changed learning objectives but maintained the structure to facilitate interannual comparisons. Results of the annual surveys guided the continued development of course materials and topics covered. In the early years of the workshop, we had consultants perform in-classroom observations and provide feedback to the instructors. Ultimately, we compiled the five years of pre- and post-survey data and performed a longitudinal analysis.

In the pre- and post-surveys, learners were asked to indicate the extent to which they understood specific learning outcomes or skills covered in the course, with ratings (e.g. Strongly Disagree, Disagree, Agree, and Strongly Agree (**Table 3**)).

**Table 3.** Representative survey questions for the “Computational Understanding” scale.

|  |
| --- |
| I know how to process Illumina data |
| I understand what per_library_stats.py does |
| I know how to run R |
| I know the main differences in analyses offered by QIIME(4) and mothur(5) |
| I am familiar with .biom(6) formatted files |
| I can name at least two different microbial metagenomic databases |
| I know what an R package(7) is |
| I understand the structure of an OTU table |
| I know what a kmer is |
| I know the difference between alpha and beta diversity |
| I know how to visualize microbial metagenomic data |
| I know how to use metadata to guide community analyses |
| I know how to assemble shotgun metagenomic data |

We also used “real-time” assessment during the workshop by replicating formative assessment strategies found to be effective in Software Carpentry workshops (8–10). Each participant was given a green (“I’m doing okay”) and a red (“I have a question”) sticky note to stick onto their open laptop during tutorials. This visual cue allowed instructors to quickly survey the classroom and determine learners’ comfort level, and to attend to any student who was struggling during tutorials. Furthermore, it allowed students to continue working through tutorials or troubleshooting without the need of raising their hand. We also employed “minute cards”. After each tutorial, students wrote what went well on the green sticky note and what could be improved on the red sticky note. Instructors and TAs read through notes during breaks to quickly identify gaps in understanding. This allowed us to identify gaps and make adjustments (e.g., in speed) in the subsequent instruction period.

### Building community resources and peer networks

We were dedicated to promote a welcoming and supportive learning environment. We presented a code of conduct in the welcome lecture so that it was clear that any questionable conduct was grounds for dismissal. We used the online “etherpad” for shared note taking to maximize engagement and inclusivity. We did our best to accommodate learners with families, providing private housing to families and learners with special requirements.

We aimed to build a peer learning community and to provide resources to support learners beyond the workshop. We offered an informal meet-and-greet on the arrival travel day and get-to-know-you lighting presentations after the first full day. These interactions allowed learners to identify peers with common research interests early in the workshop. We created a workshop website and public repository on GitHub so that learners (and outside parties) could access EDAMAME learning materials. Linked content included lectures, hands-on tutorials, and reference lists. These materials have been shared openly, with most content licensed CC-BY, so all course registrants and anyone else could have access. We also shared group email lists and encouraged social media outreach via Twitter and blogging. An EDAMAME meet-up was also held at the International Society for Microbial Ecology 2016 meeting in Montreal, CA.

### Pre- and post-survey comparisons and qualitative interviews

Ninety-seven percent of EDAMAME learners from 2014 to 2018 rated the workshop overall in the top evaluative categories, “good” to “very good.” (**Figure 3**). A comparison of pre- and post-assessment learner-reported learning gains and/or confidence with the major learning objectives of EDAMAME show gains in all sub-categories of learning reported (**Figure 4**). There were largest gains between the pre- and post-assessments with Computational Understanding (**Figure 4B**) and Perception in Ability (**Figure 4C**).

**Figure 3.**
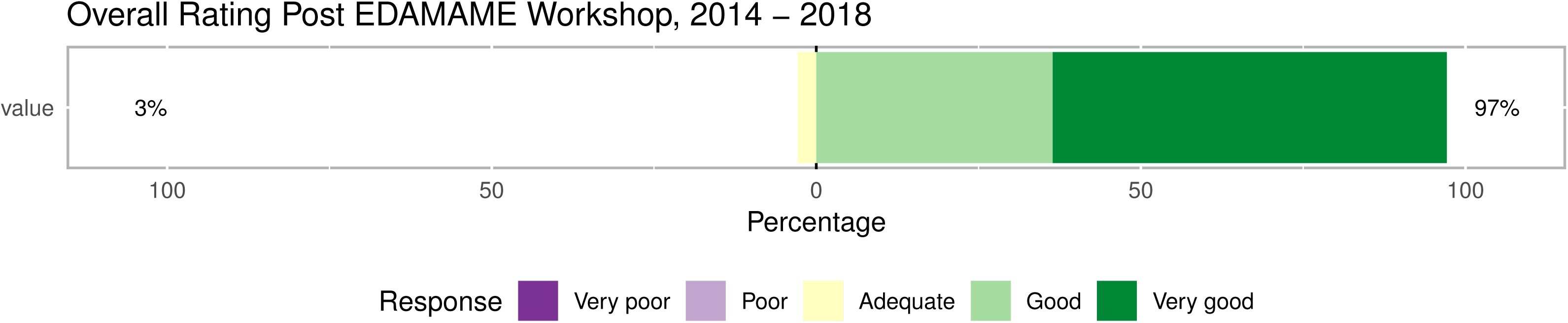
Overall EDAMAME assessment 2014-2018.

**Figure 4.**
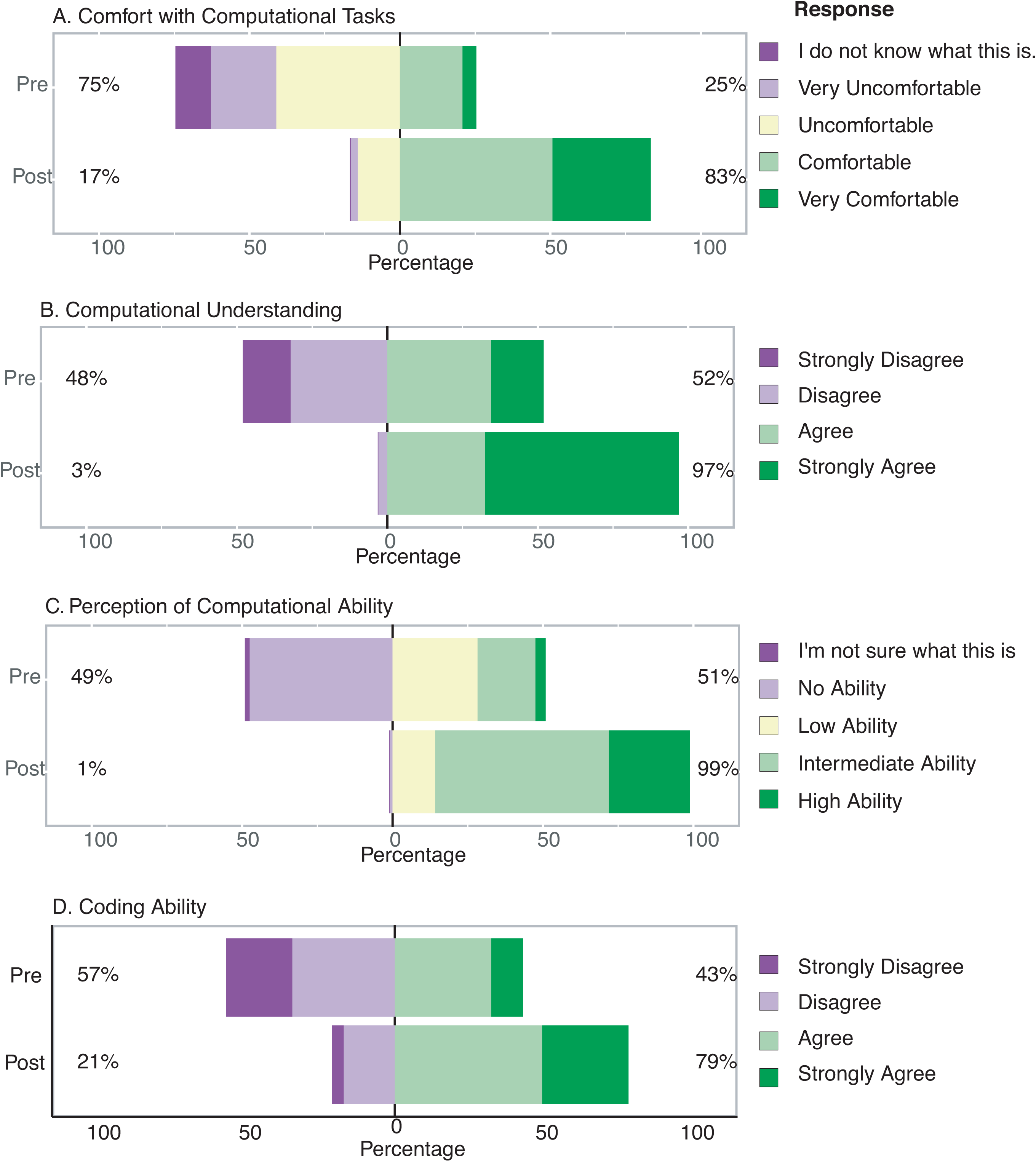
Summarized comparison of self-reported learning gains between pre- and post-workshop assessments, aggregated over 2014-2018. (A) Comfort with computational tasks; (B) Computational Understanding;(C) Perception in Computing Ability; and (D) Coding Ability.

We also asked short-answer questions at the end of the survey, in which learners were asked to design an experiment and report how they would process and analyze microbial community high-throughput sequencing data. We observed increased sophistication in the responses to the short-answer questions from the pre-to post-survey, with some learners leaving the questions blank in the pre-survey and then providing thorough answers in the post-survey. This suggests large gains especially for learners who were new to high-throughput sequence analysis.

Qualitative interviews from 9 learners who attended EDAMAME from 2014-2016 (each spending 25-40 minutes with the interviewer, **Table 4**) suggested that this group of learners were largely satisfied with the workshop and appreciated the attentiveness of the TAs and instructors as well as the red/green sticky note mechanism for soliciting help in real time. However, some of these learners also felt that there was too much material covered in the workshop and reported that they struggled to keep up with the pace of the course (“Content overwhelm”). Finally, we had many interviewed learners state that the workshop and materials covered made a positive impact on their career and research.

**Table 4.** Representative comments from interviews. The sample is small at nine attendees, but each interviewee spent somewhere between 25 and 40 minutes discussing their experience at the workshop, its impact on their professional life and walking through the agenda for their year’s workshop to give detailed feedback. While it is a small sample, each person contributed a lot of information. There were two respondents for 2014, four for 2015 and three for 2016. Each quote is labeled with the year the respondent participated in the workshop

|  |  |
| --- | --- |
| Positive overall comments | "EDAMAME was an inspiring introduction into microbiology. I thought the kind of analyses you could do with microbiology was really interesting. I really got pulled in on the data science part." (2014)<br>"It was definitely one of the most effective workshops I've been to." (2016)<br>"Very comprehensive, reached a lot of people from different backgrounds who were interested in analyzing microbial communities. I thought it provided a good survey of the tools that were available and it brought in some experts." (2014) |
| Content overwhelm comments | "I loved it, I had a blast. It was exhausting. It was a lot of fun, I learned a lot. I kind of felt overwhelmed." (2016)<br>"I appreciated the workshop for its usefulness, it's a lot to take in. We need time to process. It's nice to have a bit of a breather. For someone who was new to the field like me. I needed a bit of time to digest." (2014)<br>"It was pretty intense for me. I had never done any kind of code work before. This was really my first introduction..." (2015) |
| Career Impact Comments | "I can say that the course inspired me and put me on my path and inspired me to think about different ways to do analysis. They talked a lot about the different tools that were available." (2014) |
|  | “It was a great workshop. It really helped me in my career path. It’s opened a door for me to get into bioinformatics.” (2015)<br>“It really propelled my graduate school career and has pushed me... I took away the basic tools and I’ve been able to grow from that... I know how to make a pipeline. I know the basic structure and they gave that to me.” (2015)<br>“I’m one of the few people at [my workplace] who can analyze sequence data.” (2016) |

## Discussion and Lessons Learned

We offer suggestions from our experiences for running an effective microbiome analysis workshop (**Box 2**). EDAMAME’s content changed from 2014 to 2018 to meet changing learner needs. These changes were guided in part by the applicants’ responses to questions about their dataset and their expectations for the workshop. For example, amplicon analysis (e.g., 16S rRNA gene sequencing) was favored in early years while untargeted metagenome analysis was favored in later years. Similarly, proportionally fewer students in 2018 were novice to the command line or R, but the majority of the class appreciated the refresher. Some of the learners with self-taught experience embraced the opportunity to re-learn the “correct” approaches and to gain missing foundational knowledge. Several tutorials were popular every year. For example, there was a consistent demand for ecological statistics and “supporting” skills like GitHub/version control, and cloud computing.

**Box 2: Lessons Learned**

1. Regularly evaluate and change content to meet changing learner needs.
2. Maintain a high instructor to learner ratio.
3. Provide consistent workshop timing and fill the “middle-ground” duration needs of learners.
4. Understand the pros and cons of cloud computing for a workshop, and plan use of these resources well in advance.
5. Reach the broadest applicant pool of learners who have the potential to have the most gains from the training.
6. Consider the trade-off in workshop value (including instructor time) and maintaining economical costs to learners.
7. Plan well in advance to achieve best outcomes for applicants who require a US VISA and international travel plans to attend the workshop.
8. Almost all learning engagement needs to happen on-site; efforts to engage learners pre-workshop were ineffective.
9. Scheduled classes should teach to the majority of learners to accomplish all our learning objectives. Office hours can help struggling learners catch up.
10. A welcoming and inclusive environment creates a positive workshop experience and is essential for effective learning.

High instructor to learner ratio was essential for the success of the hands-on EDAMAME workshop. In the years that we had the lowest instructor to learner ratios (e.g., in 2014 and 2015, **Table 2**), the TAs and instructors anecdotally reported exhaustion while the learners craved more attention. In addition to formal instructors, learners could assist one another. To facilitate peer learning, we arranged the classroom in tables with groups of two or four. We also encouraged learners to support one another with troubleshooting in the time that it would take for a free instructor to come to assist

Regardless of the length of the course, several learners indicated in their post-assessments that more time at the workshop was needed each year. However, learners who were faculty or staff researchers shared (in informal conversations) that they would have been unable to commit to a longer workshop due to other professional responsibilities. We noted that there were other offerings for multi-week workshops e.g., STAMPS), as well as several one-or two-day workshops at professional society meetings and pipeline-specific training (e.g., mothur and QIIME).

Timing the workshop had several challenges. EDAMAME was held in the summer, and we tried to avoid scheduling it for the same week as major microbiology conferences, like the American Society for Microbiology Microbe meeting, the International Symposium on Microbial Ecology (ISME) and Ecological Society of America meetings. Because microbiome analysis spans multiple disciplines, it was hard to avoid all of the large conferences that microbiome researchers may attend. We also had to change the timing workshop every year to accommodate the KBS event schedule. As EDAMAME grew in popularity, some learners applied for fellowships or travel awards to support their training, but the annual change in timing made it difficult for students to plan. Moving the workshop to a dedicated conference site (e.g., a hotel) may help with consistent timing, but it would also increase the cost to learners.

We found that using cloud computing streamlined course content and democratized access. We used the Amazon Elastic Compute Cloud (EC2), which was cost effective and available to students who do not have access to high performance computers at their home institutions. In early years, we guided learners through software installation on the EC2, but in later years, we installed software in advance to focus on moving data to and from the EC2. Using the EC2 presented a challenge for learners who were affiliated with government agencies or research laboratories (e.g.,US Environmental Protection Agency, US Geological Survey) because of their need for additional security and management approval prior to installing new software or moving data. While we did not have a perfect solution for these learners, we began to anticipate their needs and prompted them in advance to receive required permissions. Another hurdle with using the EC2 was the changing way that Amazon provided student or educational computing resources over the years. In some years, Amazon provided individual credits to learners and in others required the instructors to apply for an educational grant. Cloud computing logistics needed to be anticipated about nine months in advance, but in years where individual email addresses were needed, it was impossible to prepare until after admissions were finalized, which typically occurred 4 - 6 months in advance of the workshop. We also note an issue for some international learners who did not have credit cards compatible with Amazon requirements to enroll for an EC2 account, and for these learners we had to share our own accounts or create accounts for them.

While our applicant pool and learner demographics reflected balance in gender, discipline, and academic level, EDAMAME fell short of its racial diversity goals. We could have benefitted from improvement in advertising the course to reach a broader pool to attract more applicants of color. We largely advertised on social media and through word of mouth. We recommended to specifically advertise to key target learner groups, like those underrepresented in the sciences who may be expected to have less access to the training. On a positive note, we have evidence that EDAMAME was reaching socioeconomic diversity goals, as two interview respondents were clear that they would not have had the same opportunity for training and advancement given their lower income backgrounds if it had not been for EDAMAME.

A final lesson to share is the balance between course value and learner costs. In its first years, EDAMAME was funded piece-meal by generous sponsors. We experimented with a mixed enrollment model of offering EDAMAME for university credit to local students and for fee to outside students, but many of the local students could not afford the summer tuition required for the credit hours. Then, EDAMAME was funded by external federal grants. We began to charge modest workshop fees ($325) to support items that could not be covered by the grant (e.g., coffee, snacks). As soon as we began to charge workshop fees, the majority of applicants began to request financial aid. We realized that many of the learners, mostly graduate students and post-doc, were paying for the workshop personally, so we then worked to waive fees for eligible students in need and offer scholarships for students with international travel. By contrast, the instructional team did not have enough funds to fully pay the TAs and instructors, who largely volunteered their time because they believed in the mission of the training. Guest instructors and lecturers generously volunteered their time as part of their broader impacts, and the workshop covered their travel expenses along with room and board at KBS. Thus, there is inevitable tension balancing instructor compensation and course affordability.

How much does it actually cost to run a workshop like EDAMAME? The first year, we ran the workshop for less than $14,000; students paid their own expenses of room and board; and no workshop fees were charged. This face amount did not include substantial additional support that was provided via shared logistics with the ANGUS workshop, which was occurring at the same time at Kellogg Biological Station. It also did not include any support for personnel, which was the largest true expense. Ideally, there would have been an annual budget for instructor and TA summer salaries, a logistics coordinator salary, and hourly salary for undergraduate labor during the course. We also realized that unless we could procure funds to support personnel, the training may not be valued as highly by institutions and peers, and may instead be perceived as a cost to other scholarly activities. We were grateful for the support of the NIH 2015-2018 and the USDA 2017-2018. The second biggest expense was be financial aid to offset costs of room and board and workshop fees to learners who needed it, which we provided in 2017 and 2018 to qualified learners, with USDA support. The third biggest expense was the educational consultant to evaluate the course as a neutral third-party, which was $5,500 to $6,000 per evaluation. The remaining expenses were conference services at Kellogg Biological Station, and lodging and travel expenses for the instructional team and guest speakers. In summary, there is a trade-off between the course cost, inclusive of the real value of instructor/TA time, and workshop affordability for the learners.

## Future directions

While the data indicate that EDAMAME workshop was effective, a limited number of learners can be accommodated per year, and there is high effort from the instructional team to support them. This is a low-throughput model of skill development. We are eager to reach a larger learner pool than what we could accommodate in the classroom. In 2016, we experimented with live engagement of three to five remote learners (varied by tutorial) using free conference calling and screen sharing resources. The remote learners participated as a group at the same location. They engaged with the lectures and tutorials as fully as possible (but missed out on the guest lectures and other events). This added a mild distraction for the on-site learners, but the workshop proceeded relatively smoothly. The biggest hurdle was engaging with the remote learners during tutorials, as they had no classroom support. It is possible that a remote learning workshop could be successful, given an appropriate investment into conference technology, an on-site coordinated dedicated to its logistics, and an enhanced instructional team with traveling TAs dedicated to the remote classrooms.

The content of EDAMAME remains freely available online, but parts of the content are also being transitioned to local offerings. Many universities desire more offerings of online or digitized curriculum, and there is a question of how to balance the university’s need to provide quality instruction for tuition with the open-science philosophy of providing free, democratic access to information. At Michigan State University, we are developing a graduate-level learning module on microbial metagenomics that includes amplicon and untargeted metagenome analysis pipelines. The 1-credit metagenomics module includes hands-on tutorials, is offered twice a week for one month and is accompanied by pre-recorded lectures. Post-doctoral trainees or faculty can enroll for a modest fee. Those based on EDAMAME materials, the modular content at Michigan State covers less content because there are prerequisite modules required for enrollment. Learners already have familiarity with the command line, with submitting jobs to the high-performance computing cluster, and with fundamentals of microbial genome analysis. EDAMAME materials have also been expanded to teach international workshops including, a metagenomics one-day crash course in Rio, Brazil and a one-week microbiome analysis workshop at Centro de Investigaciones Biológicas del Noroeste in La Paz, Mexico. In addition, more general tutorials (e.g., shell, GitHub, etc) remain available from other efforts, including Software and Data Carpentry, and short format 2-day workshops are available at scale through The Carpentries (http://carpentries.org) on these skills.

Finally, we seek to maximize the impact of EDAMAME by offering this kind of training to those who need it most. We hope that the impact of our trainees training others is a lasting legacy of EDAMAME. We have found that our international learners have benefited immensely from this course, as they are challenged by access to compute resources or training. Going forward, we hope to continue to identify target audiences who could both benefit from our training and extend its impact broadly. Additionally, sequence analysis will continue to evolve with technologies, impacting the depth and breadth of scientific questions and experiments that are imaginable. We hope that our course content can continue to remove obstacles for scientists who wish to engage in these technologies.

## Materials and Methods

This research was exempt under IRB ID# i052533 (standard educational practices), as reviewed by the Michigan State University Biomedical, Health Sciences Institutional Review Board (BIRB) and Social Science, Behavioral, Education Institutional Review Board (SIRB).

Data analysis for the pre- and post-survey assessment and associated reports were generated by an outside research consultants. Final reports for the years 2016, 2017 and 2018, were written by Beth M. Duckles, PhD of Insightful, LLC and for years 2014 and 2015, reports were written by Julie Libarkin of STEM ED. LLC. Code is available at [https://github.com/ShadeLab/EDAMAMESurveys]. Beth M. Duckles of Insightful also conducted qualitative interviews and provided final demographic summaries.

## Acknowledgements

We thank C. Titus Brown for encouraging us to get EDAMAME started and for sharing ANGUS resources to support our launch. We are indebted to our EDAMAME guest speakers and instructors. We express immense gratitude to every single one of our TAs for their time, enthusiasm, and commitment to EDAMAME training. We thank the EDAMAME learners for their eager participation, humor, and patience. We also thank the Ribosomal Database Project and Institute for Cyber-Enabled Research at Michigan State for sharing their talents and resources.

